# Non-structural carbohydrate concentrations in woody organs, but not leaves, of temperate and tropical tree angiosperms are independent of the ‘fast-slow’ plant economic spectrum

**DOI:** 10.1101/2021.04.20.440698

**Authors:** J.A. Ramirez, D. Craven, J.M. Posada, B. Reu, C.A. Sierra, G. Hoch, I.T. Handa, C. Messier

**Author notes:** Ramirez, J.A. and Craven, D. contributed equally to this work as first authors. Shared senior authors (I.T. Handa, C. Messier).

## Abstract

**Background and Aims:** Carbohydrate reserves play a vital role in plant survival during periods of negative carbon balance. Considering active storage of reserves, there is a trade-off between carbon allocation to growth and to reserves and defense. A resulting hypothesis is that allocation to reserves exhibits a coordinated variation with functional traits associated with the ‘fast-slow’ plant economics spectrum.

**Methods:** We tested the relationship between non-structural carbohydrates (NSC) of tree organs and functional traits using 61 angiosperm tree species from temperate and tropical forests with phylogenetic hierarchical Bayesian models.

**Key Results:** Our results provide evidence that NSC concentrations in woody organs and plant functional traits are largely decoupled, meaning that species’ resilience is unrelated to their position on the ‘fast-slow’ plant economics spectrum. In contrast, we found that variation between NSC concentrations in leaves and the fast-slow continuum was coordinated, as species with higher leaf NSC had traits values associated with resource conservative species such as lower SLA, lower Amax, and high wood density. We did not detect an influence of leaf habit on the variation of NSC concentrations in tree organs.

**Conclusions:** Efforts to predict the response of ecosystems to global change will need to integrate a suite of plant traits, such as NSC concentrations in woody organs, that are independent of the ‘fast-slow’ spectrum and that capture how species respond to a broad range of global change factors.

## INTRODUCTION

Carbon allocation to growth is a fundamental process that underpins global variation in plant functional traits, which describes quantitative differences between resource acquisitive and resource conservative species (Grime et al., 1997, Diaz et al., 2004, Wright et al., 2004, Chave et al., 2009, Reich, 2014, Díaz et al., 2016). These trade-offs reflect variation among plant traits for species that differ in growth form, size, and evolutionary history (Díaz et al., 2016, Donovan et al., 2011, Reich, 2014, Reich et al., 1999, Wright et al., 2004, Reich et al., 1997). For example, fast-growing, resource-acquisitive species, typically have high specific leaf area (SLA), high leaf nutrient concentrations, and low wood density (hereafter ‘fast’ species). In contrast, slow-growing, resource-conservative species, are characterized by low SLA, low leaf-nutrient concentrations, and high wood density (hereafter ‘slow’ species). While ‘slow’ trait values imply high construction costs, they allow trees to enhance resilience to different biotic or abiotic stress factors (Coley et al., 1985, Poorter and Kitajima, 2007).

To increase recovery and survival following periods of negative carbon balance caused by the loss of tissues that reduce the capacity to photosynthesize and take up nutrients and water, trees store and mobilize carbohydrate reserves (Atkinson et al., 2014, Canham et al., 1999, Kobe, 1997, Myers and Kitajima, 2007, O’Brien et al., 2014, Poorter and Kitajima, 2007). Thus, a significant fraction of the carbon captured by photosynthesis (between 1 and 19%) is allocated to carbohydrate reserves in the form of non-structural carbohydrates (NSC) (Hoch and Körner, 2003, Hoch et al., 2003, Landhäusser and Lieffers, 2003, Martínez-Vilalta et al., 2016, Piper et al., 2009, Würth et al., 2005). In general, NSC are comprised of low weight sugars and starch (Hoch et al., 2003). Sugars are mobilized easily and used for short-term metabolism (i.e. within a growing season), while starch is stored in a more recalcitrant form for long-term use (up to several decades (Carbone et al., 2013)) during periods of severe stress (Chapin et al., 1990, Dietze et al., 2014, Hartmann and Trumbore, 2016, Martínez-Vilalta et al., 2016).

In contrast to evergreen species that maintain photosynthetically active tissues that provide resources for plant functions all year (Fajardo et al., 2013), NSC support physiological activity during dormant periods and the flushing of new leaves of deciduous species (Fajardo et al., 2013, Gaucher et al., 2005, Gough et al., 2009, Klein et al., 2016, Messier et al., 2009, Newell et al., 2002, Würth et al., 2005). Additionally, NSC increase resilience to natural or anthropogenic disturbances providing the energy for the vital functions of plants (i.e. growth, defense, reproduction, resprouting, and survival) (Chapin et al., 1990, Kozlowski, 1992, Dietze et al., 2014, Powers, 2020). Availability of NSC may drastically reduce the risk of mortality through carbon supply to metabolism following drought periods (O’Brien et al., 2014, Doughty et al., 2015, Rowland et al., 2015, O’Brien et al., 2020, Piper and Paula, 2020, Signori-Müller et al., 2021). For instance, droughts are common in regions like the northern tropical Andes during the El Niño phenomenon (Poveda et al., 2001, Pinilla Herrera and Pinzón Correa, 2016), which usually decrease NSC concentrations in tree organs (Piper and Paula, 2020). Additionally, although the incidence of fires is uncommon in high elevation Andean forests (unless caused by anthropogenic factors), recent increases in the strength of the El Niño phenomenon has led to more burned area (Armenteras-Pascual et al., 2011), where resprouting species may require NSC as a pathway to regenerate (Poorter et al., 2010). In temperate forests of eastern North America, wind storms, ice storms, and insect outbreaks are the major natural drivers of forest dynamics (Runkle, 1982, Frelich and Lorimer, 1991). Thus, high concentrations of NSC may increase the probability of trees that restore their photosynthetic tissues and survive (Dietze et al., 2014, Hartmann and Trumbore, 2016, Martínez-Vilalta et al., 2016).

NSC concentrations increase via accumulation and reserve formation (Chapin et al., 1990). NSC accumulation is a passive process driven by the balance between photosynthesis supply and the demand of carbon for growth and respiration (Chapin et al., 1990, Sala et al., 2012, Dietze et al., 2014). Reserve formation can be also an active process in which NSC compete for carbon with plant growth and other physiological processes such as defense, even when carbon resources are limited (Chapin et al., 1990, Sala et al., 2012, Dietze et al., 2014, Weber et al., 2019). Active storage of NSC suggest an allocation-based trade-off between carbon allocated to growth and to reserves and defense (Kitajima, 1994, Kobe, 1997, Myers and Kitajima, 2007). Thus, tough leaves and dense woody organs suggest greater carbon investment in defense traits to resist and to recover from biotic and abiotic stress (Poorter and Kitajima, 2007, Poorter et al., 2010), which co-vary with carbon allocation to reserves, especially in roots (Kitajima, 1994, Myers and Kitajima, 2007). Also, since a higher SLA indicates a higher light capture potential, a higher net photosynthetic rate, and higher concentrations of foliar nutrients such as N (Wright et al., 2002, Wright et al., 2004), an increase in SLA may lead to an increase in the proportion of metabolically active carbon allocated to growth of woody organs (Shipley et al., 2006, Li et al., 2016), which may compete with storage in fast growing species. However, it remains uncertain whether the ‘fast-slow’ plant economics spectrum (Reich, 2014), which captures variation in life-history strategies, varies in coordination with NSC concentrations in leaves and woody organs.

An alternative hypothesis is that NSC concentrations are decoupled from, or are orthogonal to, the ‘fast-slow’ plant economics spectrum. This pattern would suggest that variation in NSC concentrations is uncorrelated with ‘effect’ traits, which are associated with species’ effects on ecosystem functioning, but may form part of an independent axis of ecological variation including a broader suite of ‘response’ traits whose diversity may play a role in determining the resilience of ecosystems to global change (Suding et al., 2008, Mori et al., 2013). The extent to which a trait-based spectrum of ‘resilience’ is generalizable is of basic and applied importance as it will contribute towards improving predictions of how ecosystem functioning responds to global change.

We therefore examine how key functional traits associated with the ‘fast-slow’ plant economics spectrum, as well as biome and leaf habit, vary with NSC concentrations for each tree organ in angiosperm tree species, a central issue for predicting the role of NSC in the resilience of trees that differ in life strategies. We sampled 61 angiosperm tree species across tropical and temperate biomes. We test the hypothesis that, once accounting for phylogenetic relationships among species (Freckleton et al., 2002), NSC concentrations in woody organs and leaves will be coordinated with plant functional traits that underpin the ‘fast-slow’ plant economics spectrum. Further, we anticipate that species with ‘slow’ traits associated with greater carbon investment in defense and conservative ecological strategies, such as a low SLA, high tissue density, and low concentrations of leaf nutrients will accumulate more NSC in woody organs (stem, branch and root) than species associated with acquisitive or ‘fast’ ecological strategies.

## MATERIALS AND METHODS

### Research sites

We performed this study in a deciduous temperate forest (DTF; Mont St-Hilaire, Quebec, Canada) and in an upper montane tropical forest (UMF) and a lowland tropical forest (LTF) in Colombia (Supplementary information, Table S1). These three sites were selected to generate contrasts in latitude, seasonality (temperate versus tropical), and elevation (lowland and upper montane forest (Colombia) (Fig. S1). Each study site within each biome had been protected and had not experienced recent anthropogenic disturbances (at least during the last 20 years). In the LTF, climate does not exhibit marked seasonality in terms of temperature and precipitation. The climate in the UMF exhibits a bimodal variation of precipitation between the rainy and dry seasons; the first dry period lasts from November to March, while the second one from June to August. In contrast, the climate in the DTF is characterized by strong intra-annual variation in temperature, with average sub-zero temperatures from November to March, mild and wet summers (June -September) and a growing season from May to October (Fig. S1).

### Field sampling

We sampled a total of 61 mature native tree species (see species list and leaf habit in Table S2) across the three study sites in 2012. In the temperate forest site, samples were taken after bud break and leaf maturation (May) and at the end of the growing season (October) to capture seasonal dynamics in carbon reserves that are typical of northern temperate forests. Samples in the tropical forest sites were taken between January to May, which is mostly a rainy period in the LTF and coincides with the transition from the dry to the rainy season in the UMF. We sampled before the onset of the rainy season (i.e. before leaf out) to capture the previous year’s carbon reserves. At each site, we selected abundant tree species for sampling. In Colombia, species were selected by consulting with researchers familiar with local ecosystems in order to have a representative sample of the plant communities since no biomass or abundance data were available. In Quebec, species were selected based on abundance data of Mont St. Hilaire (Maycock, 1961, Arii et al., 2005), which is why we did not sample evergreen tree species in the DTF. Botanical samples of all tropical species were verified and deposited at the Medellín Botanical Garden Herbarium Joaquin Antonio Uribe (JAUM).

At each study site, we selected the tallest trees of each species. Leaves and woody organs (branches, stems, and roots) were sampled from 3-5 individuals for each species. Current year leaves from adult plants without visible symptoms of pathogen or herbivore attack were sampled. To avoid possible effects of diurnal variation in NSC, leaf samples were collected in the early morning (Upmeyer and Koller, 1973). Leaf samples were taken from one sun-lit branch at the top of the canopy with a tree trimmer and then divided in two groups. One group was placed in paper bags for NSC measurements, while the second group was placed in plastic bags with damp tissue for measurement of leaf traits (see below). Stem samples were taken with a 4.3 mm diameter increment borer. Stem cores were taken perpendicular to the slope to reduce variability in wood density due to compression or tension. Samples of sun-lit branches 2-3 cm in diameter were obtained by cutting them with a tree trimmer. Root samples were taken with an increment borer from large surface roots ca. 50 cm away from the base of the stem. In total, we collected and analyzed samples from 326 trees.

### Non-structural carbohydrates (sugar, starch, and NSC, % of dry matter)

We placed leaf and wood samples for NSC analysis in paper bags and then in a cooler. These samples were then microwaved within 8 h after sampling to stop enzymatic activity (Popp et al., 1996). Leaf samples were ground using a ball mill and wood samples were ground using a coffee grinder with a mesh sieve. Due to the large number of samples, we selected a sub-sample of 180 samples (of a total of 1,271) using the Kennard**–**Stone algorithm (Kennard and Stone, 1969) for NSC analysis following (Hoch et al., 2002) based on variation in near-infrared spectra. Ground plant material was dissolved for 30 min in distilled water. Starch and sucrose were disaggregated in glucose and in glucose and fructose, respectively, with Clarase (*Aspergillus oryzae*, Enzyme Solutions Pty Ltd, Crydon South, Victoria, Australia) by incubation at 40°C for 15 h.

Phosphoglucose-isomerase was added to the solution and then the total amount of glucose (corresponding to total NSC) was quantified photo-metrically in a microplate photometer at 340 nm (Thermo Fisher Scientific, Waltham, USA) after conversion of glucose to gluconate-6-phosphate (hexokinase; Sigma-Aldrich, St. Louis, MO, USA). An aliquot of the original extract was treated with invertase and phosphoglucose-isomerase (both Sigma-Aldrich) to determine the amount of glucose, fructose, and sucrose using a glucose test. Starch was calculated as NSC minus sugar. We used pure starch and solutions of glucose, fructose, sucrose, and plant powder (orchard leaves; Leco, St. Joseph, MI, USA) as standards and to control reproducibility of the extraction.

Using the same dataset, we extrapolated NSC values from the 180 sub-samples to all samples using near-infrared reflectance spectra (Ramirez et al., 2015). Reflectance spectra were measured using a FT-NIR spectrometer Analyzer (Bruker MPA Multi-Purpose FT-NIR Analyzer, Bruker Optik GmbH, Ettlingen, Germany) for all samples. The reflectance spectra were taken from 800 nm to 2780 nm with a mean spectral resolution of 1.7 nm on five scans per sample. The spectral data were recorded as absorbance (log (1/R), where R = reflectance). Then, we fitted regression models that predict NSC concentrations in different tree organs (leaves, stems, branches, and roots) from near-infrared reflectance spectra using partial least squares regression and competitive adaptive re-weighted sampling (Li et al., 2009). Across all tree organs, the model fit for NSC was *r*^2^ = 0.91 (Ramirez et al., 2015). The NSC concentrations are reported here as the percentage of dry matter.

Because tree height may influence NSC allocation patterns (Sala and Hoch, 2009, Genet et al., 2009, Piper and Fajardo, 2011, Woodruff and Meinzer, 2011), we measured tree height of all sampled individuals (H, m). All sampled trees were at or close to their maximum height as they were sampled in mature forests. Tree height was measured with a TruPulse 360 laser with a resolution of 10 cm for linear lengths (Laser Technology, Inc., CO, USA).

### Functional traits

We measured 11 traits that are associated with important ecological strategies for tree functioning, productivity, and survival (Table S3) following standard protocols (Pérez-Harguindeguy et al., 2013).

#### Leaf size (LS, mm^2^), leaf thickness (LT, mm), leaf dry matter content (LDMC, mg g^-1^), and specific leaf area (SLA, mm^2^ mg^-1^)

Eight completely expanded leaves were randomly collected from the leaves of a sampled branch of each individual. Leaves were placed in plastic bags in the field with damp paper to maintain humidity. After determining fresh leaf mass, we dried leaf samples in an oven at 60 °C until constant weight. LS was measured using WinFolia (*Regent Instruments, Toronto, Canada*). LT was measured on fresh leaves as the mean of four measurements with a digital micrometer (Mitutoyo Instruments, Singapore). LDMC was calculated as leaf dry mass divided by its fresh saturated mass. SLA was calculated as the area of the fresh lamina surface divided by its dry mass.

#### Photosynthetic capacity by mass (Amass, nmol CO_2_ g^-1^ s^-1^)

Photosynthetic capacity was measured on six leaves from two sun-lit branches in both tropical forest sites using a LI-6400 portable photosynthetic system (LI-COR, Lincoln, NE, USA). Branches for photosynthesis determination were placed in a bucket of water during the measurements to avoid disruption of water transport within the xylem (Verryckt et al., 2020). The photosynthetic capacity under saturating light (A_max_) was measured at 2000 µmol m^-2^ s^-1^. Measurements were carried out under constant CO_2_ concentration (390 ppm) and leaf temperature (set at 20 °C). Leaves were allowed to acclimate to 1000 µmol m^-2^ s^-1^ and then 2000 µmol m^-2^ s^-1^ for several minutes before measurements. Photosynthetic capacity per leaf dry mass was calculated as the product of A_max_ (*μmol CO*_*2*_ *m*^*-2*^ *s*^*-1*^) and *SLA*^-1^. Photosynthetic data for tree species in the DTF were obtained from (Marino et al., 2010), which were measured in a similar manner as described above.

#### Leaf nutrients (leaf N, % and leaf Ca, leaf Mg, and leaf P, mg kg^-1^)

About 20 g of leaf tissue per tree were dried and ground to a fine powder using a ball mill. Nitrogen concentrations were determined for all leaf samples with a CN elemental analyzer (Vario MAX, Elementar, Germany). Determination of Ca, Mg, and P was performed on 100 samples using the acid digest method (Allen et al., 1974), and these results were extrapolated to all leaf samples using FT-NIR reflectance spectroscopy as described for NSC (Ramirez et al., 2015). Model fit for leaf nutrients was (*r*^2^) 0.93, 0.76 and 0.78 for Ca, Mg, and P, respectively (Fig. S2).

#### Wood density (stem density (SD) and branch density (BD), mg mm^-3^)

Samples of stems and branches were placed in plastic bags in the field with damp paper to maintain humidity, and then were soaked in water in the lab for 48 hours. Fresh wood volume was measured without bark by water displacement, and wood mass was determined after drying samples at 60 **°**C, and then again at 100 °C, to a constant weight (Williamson and Wiemann, 2010).

### Statistical analysis

#### Phylogeny

We used an updated version of the molecular phylogeny from Zanne *et al*. (2014) and Qian and Jin (2016) to build a phylogeny with the *congeneric*.*merge* function in the ‘*pez*’ R package (Pearse et al., 2015), conservatively binding species into the backbone using dating information from congeners in the tree.

#### ‘Fast-slow’spectrum

Using gap-filled, individual-level data for all functional traits, we performed a principal components analysis using the PCA function in the R package “*FactoMineR*” (Lê et al., 2008) to represent the *‘*fast-slow*’* spectrum of plant form and function (Díaz et al., 2016). Prior to principal components analysis, leaf area, leaf thickness, SLA, Amass, and leaf Ca were natural log transformed to meet normality assumptions. Because the first two axes of the PCA explain a considerable amount of trait variation (57.4%), we decided to use both in subsequent analyses (see below) and hereon refer to them as fast-slow PC1 and fast-slow PC2. The PCA suggested that DTF species, all of which are deciduous, exhibit a restricted trait variation as compared to the trait space of the other species. For this reason, we included interactions between the first two axes of the ‘fast-slow’ spectrum with leaf habit and biome in the initial models described below.

### Data analysis

#### Variance partitioning

To determine the relative contributions of biome, leaf habit, and species to variation in NSC for each tree organ, we fitted an intercept only linear mixed-effects model with a nested random effects structure (∼1|Biome/Leaf habit/Species/Tree) using restricted maximum likelihood (REML) with the *lme* function in the R package “*nlme*” (Pinheiro et al., 2020). Variance partitioning was estimated using the *varcomp* function. The variance partitions represent the amount of variation within each level, i.e. the variance partition for “species” represents interspecific variation (Messier et al., 2010). Note that variation within the tree level also includes residual variation, meaning that this partition captures intra-specific variation and error.

#### Phylogenetic signal

We estimated the phylogenetic signal of NSC concentrations of tree organs as lambda using the *phylosig* function in the R package “*phytools*” (Revell, 2012), incorporating sampling error following Ives, Midford and Garland (2007) to account for multiple observations per species. Lambda values close to 0 indicate no phylogenetic signal while values close to 1 indicate trait evolution according to the Brownian motion evolutionary model (Molina-Venegas and Rodríguez, 2017).

#### Phylogenetic hierarchical models

Because phylogenetically closely related species are likely to share similar trait values (Felsenstein, 1985, Freckleton et al., 2002), not accounting for phylogenetic relationships may reduce estimation accuracy and increase type I error rates (Li and Ives, 2017). Moreover, accounting for phylogenetic relationships in our analyses allow for direct comparisons across tree organs because different species were sampled for NSC across tree organs, and the phylogenetic signal of NSC varied markedly across tree organs (see Results). We therefore fitted separate phylogenetic multi-level Bayesian models to examine variation in root, stem, branch, and leaf NSC as a function of biome, leaf habit, the ‘fast-slow’ spectrum (fast-slow PC 1, fast-slow PC 2), and two-way interactions between biome, leaf habit, and both axes of the ‘fast-slow’ spectrum. Because we did not sample evergreen species in the deciduous temperate forest, we did not fit models with an interaction between biome and leaf habit. All interactions were initially included in all models; if the 95% credible intervals of interactions overlapped with zero, models were then re-fit without these interactions.

As sampled trees were likely to be at or close to their species’ maximum height, we expected that the influence of tree height on NSC concentrations is similar across species. However, to account for the positive correlation between tree height and NSC concentrations within species (Sala and Hoch, 2009, Woodruff and Meinzer, 2011), we included tree height as a random slope in all models. To account for phylogenetic correlations among species, we included two random intercept terms for species: one term that models phylogenetic covariance and another term that accounts for repeated measurements and other effects that may be independent of phylogenetic relationships among species (Ives, 2018). The random effect structure allowed slope and intercept parameters to vary for each species. As NSC concentrations for roots, stems, branches, and leaves were not measured on all individuals, models were fit to subsets of data for each plant organ.

We fit all models using weakly informative priors, four chains, and 1,500 burn-in samples per chain, after which 4,500 samples per chain (total post-warmup samples = 18,000) were used to calculate posterior distributions of model parameters. To reduce the number of divergent transitions, we set the ‘*adapt_delta*’ parameter within the ‘*brms*’ function to 0.99 for all models (Bürkner, 2017). All fixed effects were standardized using a z-transformation to enable comparisons across models. Model convergence was evaluated visually and by estimating ‘*Rhat*’ using the ‘*rhat*’ function, where values considerably greater than 1 indicate that models have failed to converge. We selected the distribution family by fitting each model with a Gaussian or log-normal distribution, and assessed which fit better by comparing observed data to simulated data from the posterior predictive distribution (Fig. S3). Additionally, we estimated a Bayesian *r*^2^ using the ‘*bayes_R2*’ function for each model to represent an estimate of the proportion of variation explained for new data.

As NSC from fall sampling may be a better overall indicator of current year growth-storage trade-offs in the Northern Hemisphere (Martínez-Vilalta et al., 2016), we used the fall NSC concentrations sampled in the deciduous temperate forest for all models. As a sensitivity test, we also examined if trait-NSC relationships varied across seasons within the DTF using a similar model structure as described above. All analyses were performed in R version 3.6.1 (R Core Team 2020).

## RESULTS

### Traits and NSC concentrations across biomes and leaf habit

Within each biome and leaf habit, tree species of the tropical biomes exhibited a broad range of variation in ecological strategies, in contrast to the DTF species, all of which are deciduous (Fig. 1 see Fig. S4 for individual functional traits). The first two axes of the PCA captured a total of 58.8% of variation among the 11 functional traits, the first axis capturing 34.3% of variation and the second capturing 23.1%. The first PCA axis represents traits associated with mechanical strength, defense and resource acquisition, from ‘slow’ species with high branch and stem density and LDMC, to fast species with high Amass, leaf N, and leaf P. The second PCA axis represents traits related to resource acquisition and defense, from ‘slow’ species with high leaf thickness, leaf Mg and leaf area to fast species with high SLA and Amass (Fig. 1).

**Fig. 1.**
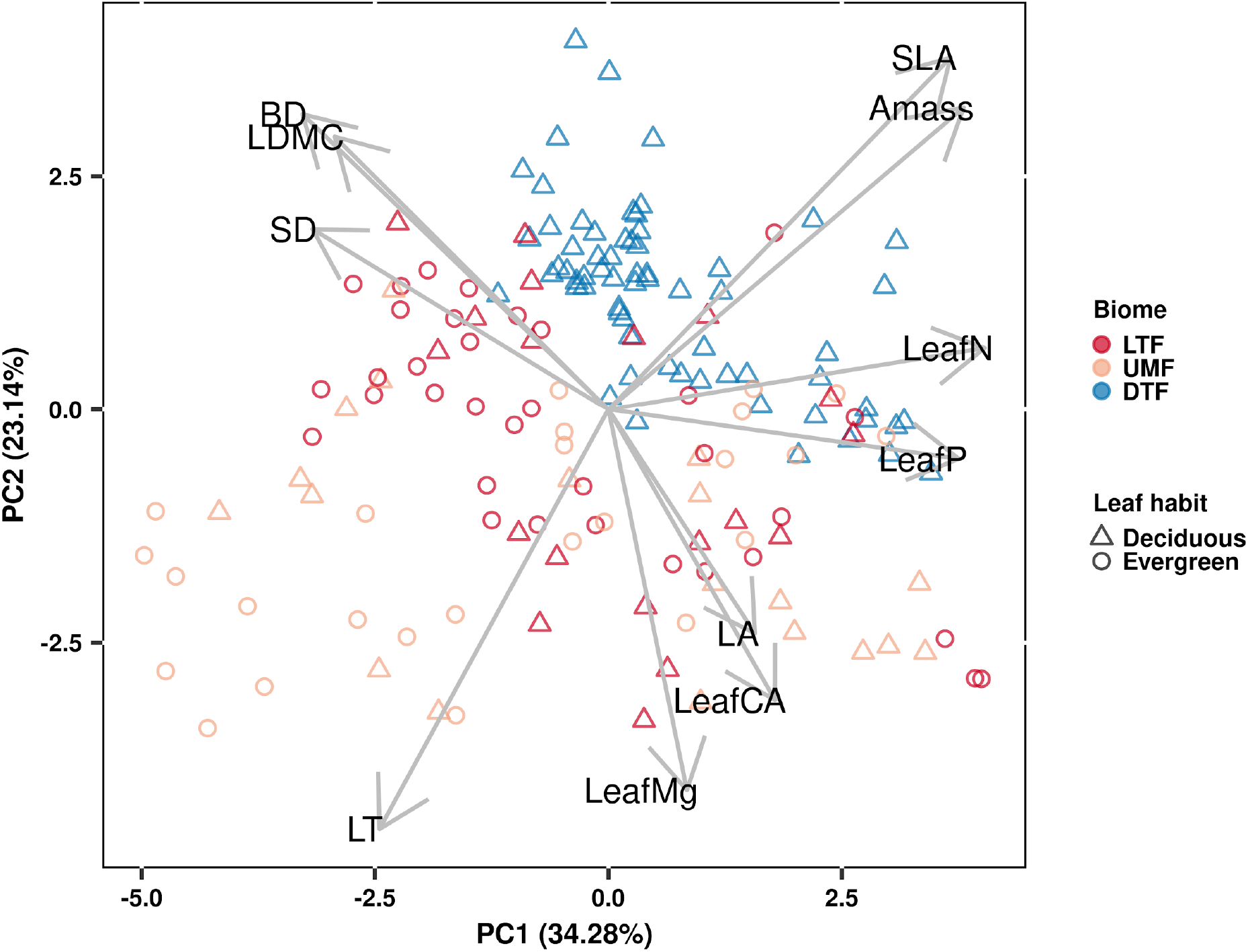
Principal components analysis of plant functional traits across biomes (*n* = 61 species). LTF: lowland tropical rainforest, UMF: upper montane forest, and DTF: deciduous temperate forest. Table S3 shows trait abbreviations.

Our analyses showed that NSC concentrations were similar across biomes for all tree organs except leaves, which were significantly higher in the DTF. However, NSC concentrations among organs were weakly correlated (*r* < 0.4, Fig. S5). Variance partitioning analysis showed that most of the variation in NSC in tree organs is explained by interspecific variation for roots, stems and branches and by among-biome variation for leaves (Fig. 2). There was a minimal influence of leaf habit on the variation of NSC concentrations for any tree organ (Fig. S6). We found broadly similar results when examining NSC concentrations in tree organs across seasons in the deciduous temperate forest (Fig. S7); only NSC in branches exhibited a higher concentration in the fall than in the spring (Fig. S7c).

**Fig. 2.**
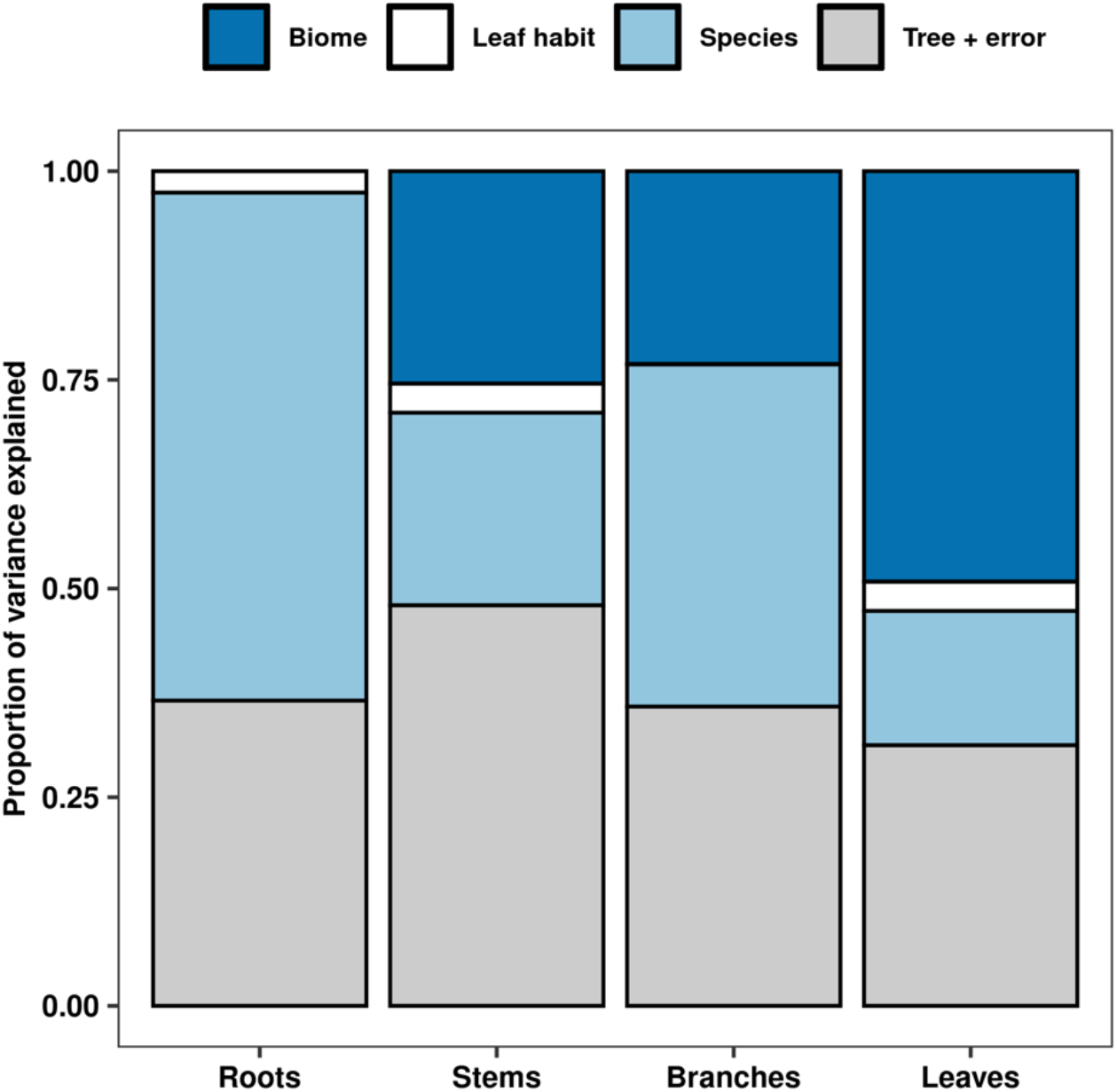
Variance partitions of NSC concentrations for each tree organ across biome, leaf habit, and species. Variation within tree + error includes residual variation, meaning that this partition captures intra-specific variation and error.

The phylogenetic signal of NSC concentrations was greater than 0 for leaves, branches and stems, indicating a moderate amount of phylogenetic signal, but not as much as would be expected under Brownian movement (Table 1). In contrast, the phylogenetic signal of NSC concentrations for roots was close to zero (Table 1).

**Table 1.** Model fit and phylogenetic signal in NSC concentrations of organs of tropical and temperate tree species. Model convergence (Rhat) and Pagel’s lambda were estimated directly by phylogenetic multi-level Bayesian models. 95% credible intervals are in parentheses.

| Organ | $r^2$ | Pagel's lambda | Rhat |
| --- | --- | --- | --- |
| Leaves | 0.66 (0.57 – 0.72) | 0.54 | 1.00014 (1.00011-1.00018) |
| Branches | 0.53 (0.38 – 0.65) | 0.12 | 1.00023 (1.00015-1.00030) |
| Stems | 0.54 (0.34 – 0.70) | 0.30 | 1.00016 (1.00011-1.00021) |
| Roots | 0.72 (0.60 – 0.81) | 0.00 | 1.00063 (1.00047-1.00079) |

### Relationships between carbohydrate concentrations and the fast-slow continuum across biomes and leaf habit

Our phylogenetic multi-level Bayesian models that examine variation in root, stem, branch, and leaf NSC as a function of biome, leaf habit, and the traits of the ‘fast-slow’ spectrum (fast-slow PC 1, fast-slow PC 2), explained a large amount of variation in NSC concentrations, ranging from 53% to 72% across tree organs (mean Bayesian *r*^2^; Table 1) and estimated NSC concentrations (Fig. 3).

**Fig. 3.**
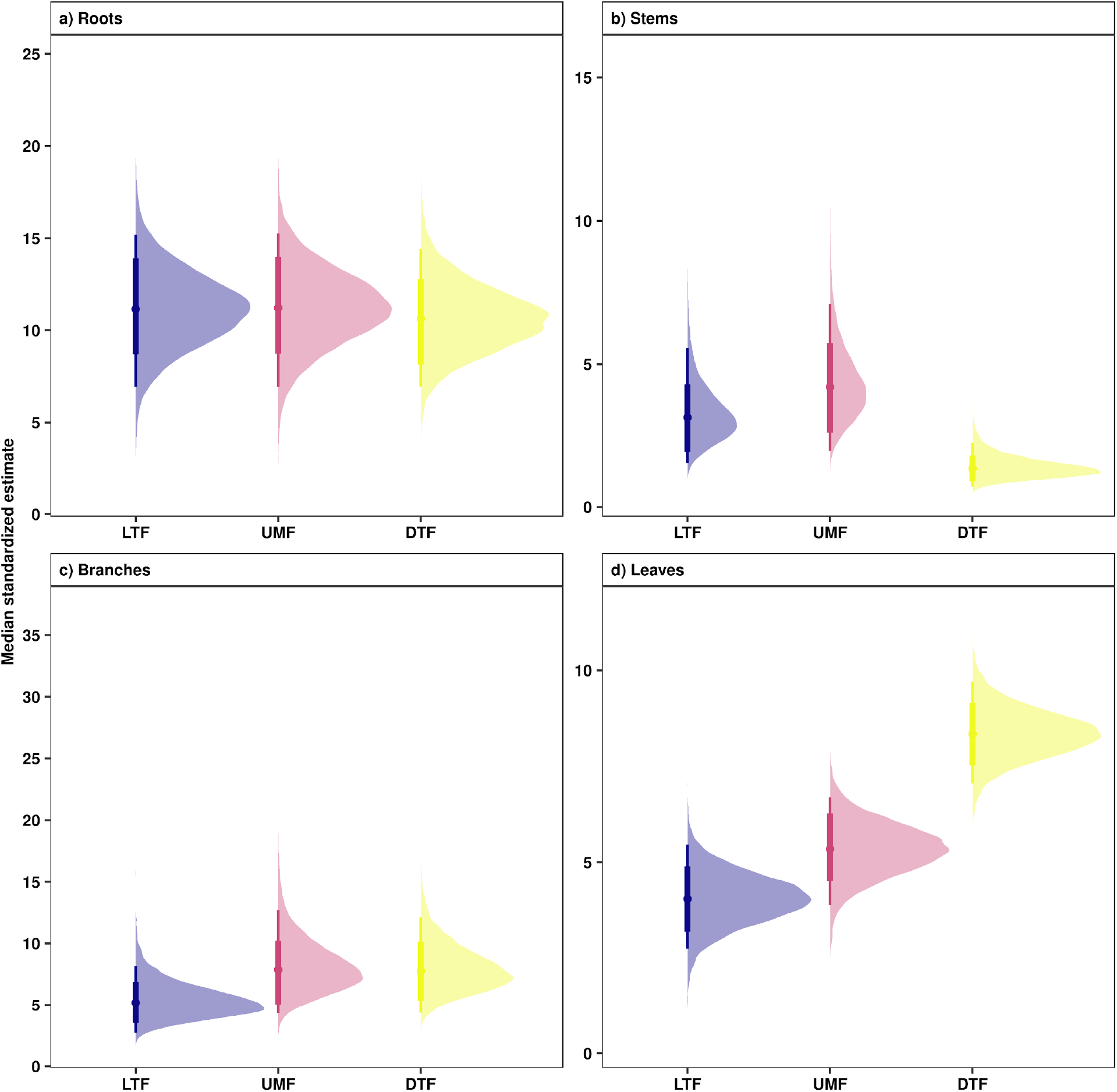
Estimated fall NSC concentrations in **a**) root, **b**) stem, **c**) branch, and **d**) leaves across biomes. Points are medians and whisker bars are 80% and 95% credible intervals and were estimated using phylogenetic hierarchical Bayesian models. LTF: lowland tropical rainforest, UMF: upper montane forest, and DTF: deciduous temperate forest.

Contrary to our expectations, the two dimensions of the ‘fast-slow’ spectrum (fast-slow PC1, fast-slow PC2) did not consistently predict variation in NSC concentrations of woody organs across biomes (Fig. 4 Table S4). Only fast-slow PC2 showed a marginally positive relationship with NSC concentrations in roots (80% credible intervals, Fig. 4a), indicating that more resource acquisitive species tend to have a higher concentration of reserves in roots. Fast-slow PC1 varied negatively with leaf NSC concentrations (Fig. 4d Table S4), indicating that resource conservative species have higher concentrations of NSC in leaves.

**Fig. 4.**
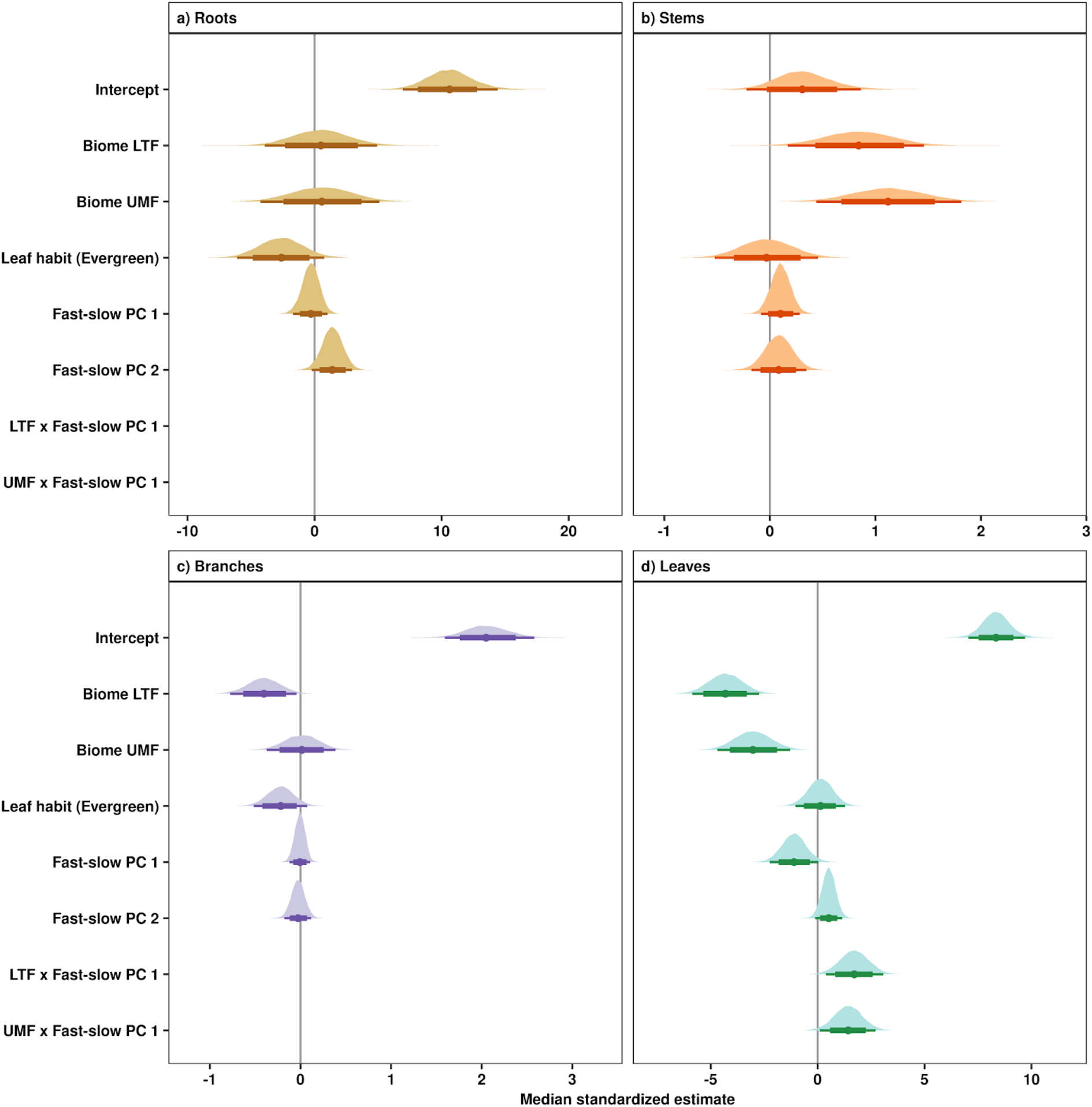
Influence of the fast-slow continuum, biomes and leaf habit on NSC concentrations of **a**) roots, **b**) stems, **c**) branches, and **d**) leaves. Points are medians and whisker bars are 80% and 95% credible intervals estimated using phylogenetic hierarchical Bayesian models. Continuous variables were z-transformed prior to analysis to facilitate comparisons (within and across tree organs). Fast-slow PC1 and fast-slow PC2 are the first two axes of a principal component analysis of fast-slow plant functional traits. LTF: lowland tropical rainforest, UMF: upper montane forest, and DTF: deciduous temperate forest. Table S3 shows trait abbreviations.

Our analysis further showed context dependent effects of the ‘fast-slow’ spectrum on NSC concentrations in leaves (Figs. 4d 5 and S8). Leaf NSC concentrations varied slightly along the fast-slow PC1 in the tropical biomes, while in the DTF leaf varied negatively with the fast-slow PC1 (Fig. 5). This result indicates that in the DTF ‘slow’ species along the first fast-slow dimension (PC1) have higher leaf NSC concentrations than fast species.

**Fig. 5.**
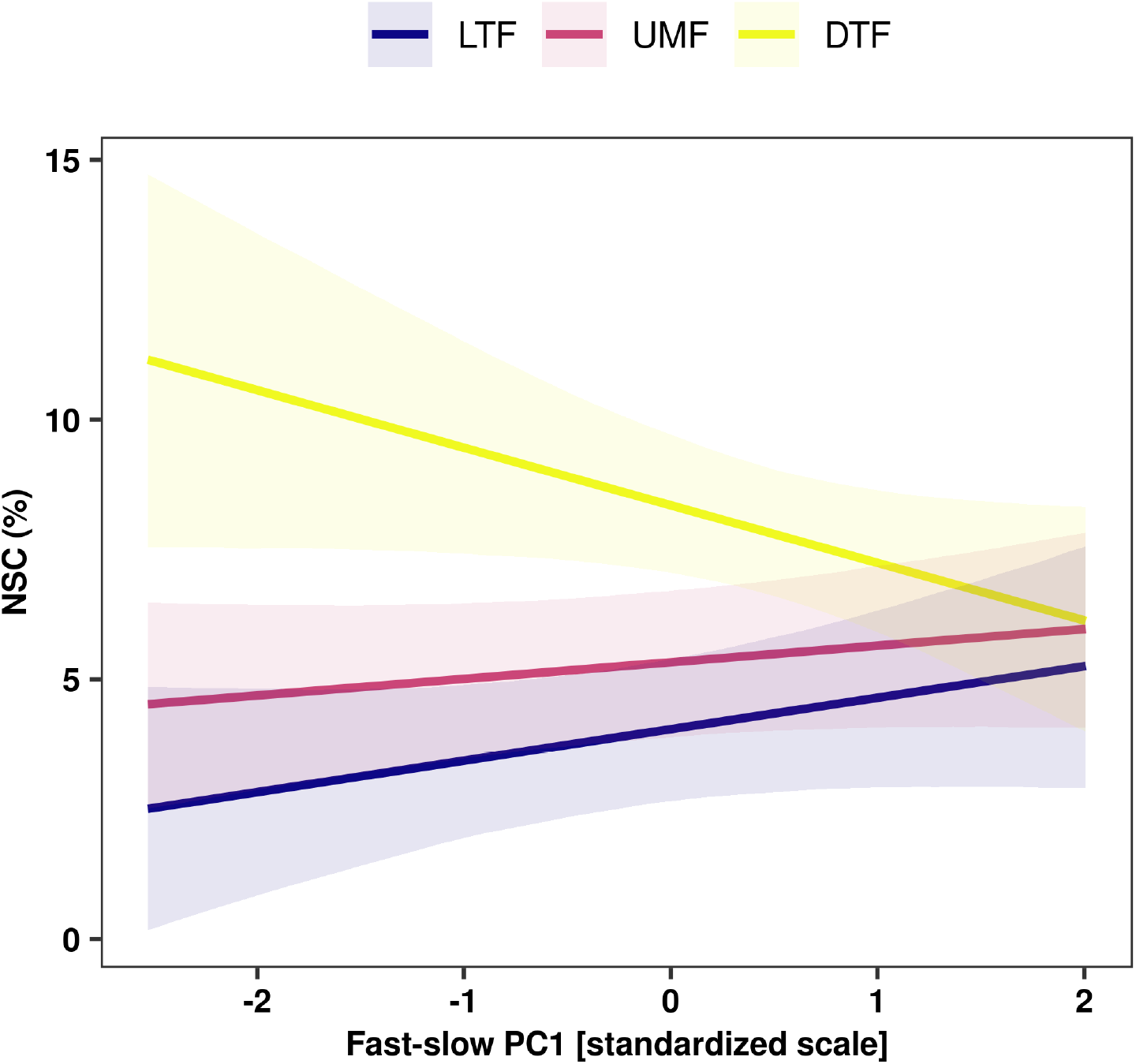
Interactive effects of the fast-slow PC 1 and biome on NSC concentrations of leaves. Solid lines are predicted fitted values using phylogenetic hierarchical Bayesian models and shaded regions represent 95% credible intervals of fitted values. Plant functional traits were z-transformed prior to analysis. LTF: lowland tropical rainforest, UMF: upper montane forest, and DTF: deciduous temperate fores.

## DISCUSSION

Our examination of the relationships between carbon reserve concentrations and functional traits of temperate and tropical angiosperm tree species revealed that variation between traits and NSC concentrations in woody organs was largely decoupled. Conversely, we found that leaves exhibited coordinated variation between NSC concentrations and traits (fast-slow PC1), which varied in direction and strength among biomes (Fig. 4 and 5).

### Coordination between carbohydrate concentrations and functional traits

In general, our results show that relationships between functional traits and carbohydrate concentrations in woody organs were not coordinated (Fig. 4). This suggests that the position of species along the ‘fast-slow’ plant economics spectrum is not predictive of NSC concentrations in woody organs, extending the findings from previous studies showing no trade-off between NSC concentrations and carbon investment (Lusk and Piper, 2007, Piper et al., 2009, Imaji and Seiwa, 2010, Piper, 2015) by considering species from multiple biomes.

A possible explanation for the lack of a consistent relationship between functional traits and NSC concentrations in woody tree organs may be that long-term allocation of carbohydrates to storage in stems or roots can take several growing seasons or years (Carbone et al., 2013, Richardson et al., 2015, Hartmann and Trumbore, 2016, Muhr et al., 2016), depending on the distance and the osmotic gradient between carbohydrate sources and sinks (Lacointe, 2000, Le Roux et al., 2001). If other trait values capture the current abiotic and biotic conditions to a greater extent than those present during the accumulation of NSC, the strength of their association with NSC concentrations may weaken with increasing age of stored NSC. Therefore, the age difference between NSC of leaves and woody organs may explain why variation in functional traits is largely decoupled from NSC of woody organs, but not from leaf NSC.

Among the studied traits included in fast-slow dimensions, it was surprising that variation in NSC in woody tissues was decoupled from wood density of stems and branches (SD and BD). Wood density has been suggested to be a proxy for both the amount of parenchyma (Ziemińska et al., 2015, Morris et al., 2016), and NSC concentrations (Plavcová and Jansen, 2015, Plavcová et al., 2016). However, parenchyma cells have multiple functional roles, e.g., acting as a water reservoir and contributing to different mechanical properties of wood (i.e., elasticity) that are independent of wood density and NSC concentrations (Ziemińska et al., 2015). Additionally, xylem structure – in order to enhance mechanical stability – places strong constraints on the storage capacity of tree stems (Plavcová et al., 2019), the dimension with which SD and BD are most strongly associated, and was decoupled from NSC concentrations in these tissues (Fig. 4).

In contrast with NSC concentrations of woody organs, we found evidence of coordination between leaf NSC concentrations and the fast-slow PC1 (Fig. 4d Table S4). The decrease in leaf NSC with increasing leaf N, leaf P, SLA and Amax and the increase in leaf NSC with increasing SD and LT suggest that species with ‘slow’ ecological strategies accumulate more NSC in their leaves than those with ‘fast’ ecological strategies. Our results therefore suggest that acquisitive trait values of leaf N, leaf P, SLA and Amax may be associated with structural support traits such as leaf toughness and leaf lifespan (Osnas et al., 2018). Even though leaf N and P are strongly correlated with Amax (Reich and Schoettle, 1988), because physiologically they play a fundamental role in both photosynthesis and starch and sucrose synthesis (Rychter et al., 2016), they may not favor NSC accumulation in temperate and tropical biomes.

Thus, our results did not fully support our second hypothesis that species with ‘slow’ traits associated with greater carbon investment in defense and conservative ecological strategies accumulate more NSC in woody organs than species associated with acquisitive or ‘fast’ ecological strategies. Our results show that species with higher NSC concentrations in roots exhibit a weak, positive tendency (80% credible intervals) to have trait values associated with ‘fast’ ecological strategies. This finding may suggest that ‘fast’ species may allocate more carbohydrates to roots than ‘slow’ species as part of their response to mechanical damage to aboveground plant organs, which may enable them to persist in areas subjected to frequent disturbances, such as winds, low intensity fires (Poorter et al., 2010, Clarke et al., 2013), ice storms (Proulx and Greene, 2001), or in human-dominated ecosystems (Uhl, 1987, Jakovac et al., 2015). We found contrasting trends in trait-NSC relationships among biomes for leaves, where leaf NSC did not vary or increase with the fast-slow PC1 in the tropical biomes but decreased with increasing the fast-slow PC1 in the DTF (Fig. 5). Several studies on woody plants have reported contrasting patterns of plant functional strategies among biomes, which have been associated with phylogenetic constraints, or selective biogeographic processes, such as adaptation to different climatic regimes or physical barriers that generate different selective pressures within communities (Wright et al., 2005, Heberling and Fridley, 2012, Heberling and Fridley, 2013). Additionally, other abiotic factors, such as soil fertility and water availability, may mediate the growth-storage trade-off at the level of NSCs by either facilitating or constraining tree growth (Breugel et al., 2011).

### Patterns of NSC storage across tree organs

While plants may remobilize nutrients and reserves between organs according to fluctuating resource availability (Maillard et al., 2015), our results show that trees accumulate large amounts of carbohydrate reserves over time in woody organs – especially in roots – regardless of their ecological strategy or species leaf habit (Fig. 2). The high NSC concentrations in roots observed in this study may indicate that roots serve as a long-term reservoir for responding to future disturbances (Clark and Clark, 1991, Poorter et al., 2010, Clarke et al., 2013), ensuring that resources are available for resprouting or leaf flush (Wiley, 2013). For example, different studies have shown that trees have sufficient reserves to rebuild the entire leaf canopy up to four times (Hoch et al., 2003, Körner, 2003, Würth et al., 2005), or even provide the carbon necessary for stem growth for up to 30 years (Klein et al., 2016). Thus, NSC stored in stems and roots probably remain stable or increase gradually over time, at least until a severe disturbance triggers an imbalance between carbon sources and sinks and initiates mobilization of reserves (e.g. Morales et al., 2020). The stability of root and stem NSC reserves is likely in contrast to the more labile, more recent NSC reserves stored in leaves and branches that support daily metabolism and annual growth (Martínez-Vilalta et al., 2016). We found lower branch NSC concentrations in the spring than in the fall of the deciduous temperate forest (Fig. S7), which suggests that carbohydrates used for leaf flush were supplied from the closest sources of reserves (branches) and not from the more distant ones (stems and roots) (Hoch, 2015, Klein et al., 2016).

Finally, NSC concentrations vary with the time of year, especially in trees subjected to strong seasonality, such as in temperate regions where carbohydrate reserves are necessary to survive the winter (Canham et al., 1999, Barbaroux et al., 2003, Li et al., 2002, Hoch and Körner, 2003, Körner, 2003, Genet et al., 2009, Carbone et al., 2013, Martínez-Vilalta et al., 2016). Thus, tree species with contrasting ecological strategies may have similar fall carbohydrate concentrations but different spring concentrations (Hoch et al., 2003, Martínez-Vilalta et al., 2016). Here, we present results based on fall NSC concentrations in a deciduous temperate forest, since they may be a better indicator of current year growth-storage trade-offs in the Northern Hemisphere (Martínez-Vilalta et al., 2016). We also evaluated NSC concentrations based on mean NSC concentrations across seasons (spring and fall in the deciduous temperate forests) to examine seasonal remobilization of carbohydrates, and to test the sensitivity of our results to seasonal variation in NSC concentrations. We found broadly similar results when examining trait -NSC relationships across seasons in the deciduous temperate forest; only fast-slow dimensions and NSC in stem organs exhibited contrasting slopes across seasons (Figs. S7 and S8).

## Conclusions

Our study tested the hypothesis that NSC concentrations in tree organs are associated with the ‘fast-slow’ spectrum of leaf and wood functional traits across biomes. In woody organs, we found that variation between NSC concentrations and functional traits were mostly decoupled, yet were strongly coordinated in leaves. Considering the concentrations of NSC in woody organs as a proxy for species’ capacity to respond to disturbances, our results imply that variation in species’ NSC concentrations is weakly associated with functional trait spectra that describe global variation in plant ecological strategies. Consequently, efforts to predict the response of ecosystems to global change will need to integrate a suite of plant traits, such as NSC concentrations, that are independent of the ‘fast-slow’ spectrum and that capture species’ resilience to a broad range of potential drivers.

## ACKNOWLEDGEMENTS

The authors thank Milena Molina, Juan Carlos Medina, Luis Carlos Galeano, David Andres Herrera, Sergio Martinez, and Mathieu Messier for their help. We also thank Alfredo Navas (Hacienda Sabaneta Natural Reserve) and Alvaro Cogollo (Medellin Botanical Garden) for their logistical support, and Daijiang Li for his input on data analysis. This research was supported by the NSERC/Hydro-Quebec research chair on tree growth control and by a scholarship from the Quebec research fund for nature and technology. DC received support from Chile’s National Fund for Scientific and Technological Development (FONDECYT No. 1201347).

## AUTHOR CONTRIBUTIONS

JAR, DC, JP, ITH, and CM contributed to the design of the research. Fieldwork was carried out by JAR. Laboratory analysis of sugars and chemical elements were performed by JAR with the support of GH, BR, and CS. JAR and DC analyzed the data. JAR led the writing of the manuscript with substantial input from DC and ITH. All authors contributed critically to improve drafts and gave final approval for publication.

