## Supplemental material for "Non-structural carbohydrate concentrations in woody organs, but not leaves, of temperate and tropical tree angiosperms are independent of the ‘fast-slow’ plant economic spectrum"

**Supplementary information**

**Table S1**. Main characteristics of the study sites.

| | Rio Claro Reserve Antioquia, Colombia (LTF biome) | Hacienda Sabaneta Nature Reserve Cundinamarca, Colombia (UMF biome) | Gault Nature Reserve Mont-Saint-Hilaire Quebec, Canada (DTF biome) |
|---|---|---|---|
| **Biome** | Lowland tropical rainforest | Upper montane forest | Deciduous temperate forest |
| **Latitude** | 5°54’04’’ N | 4°32’30’’ N | 45°32’31’’ N |
| **Longitude** | 74°51’24’’ W | 74°15’18’’ W | 73°09’11’’ W |
| **Altitude (range, m asl)** | 250 – 750 | 2500 – 3300 | 200 – 400 |
| **Mean annual precipitation (mm)** | 3882† | 1956† | 967‡ |
| **Precipitation regime** | Bimodal | Bimodal | Unimodal |
| **Mean annual temperature (°C)** | 26 | 12 | 6 (16*) |
| **Mean annual freeze-free days** | 0 | 0 | 140‡ |
| **Soil order** | Entisol | Andisol | Chernozemic |
| **Number of species studied** | 32 | 27 | 21 |
| **Disturbance history** | Reserve protected since 1976 | Reserve protected since 2008 | Reserve protected since 1958 |

* Mean growing season temperature.
† IDEAM (Institute of Hydrology, Meteorology and Environmental Studies of Colombia).
‡ Environment and natural resources. Canada.

**Table S2**. List of tree species sampled in Colombia and Canada along with its leaf habit. UMF: upper montane tropical forest, LTF: lowland tropical rainforest, and DTF: deciduous temperate forest.

| Species | Family | Biome | Leaf habit |
|---|---|---|---|
| *Acer pensylvanicum* | Sapindaceae | DTF | Deciduous |
| *Acer rubrum* | Sapindaceae | DTF | Deciduous |
| *Acer saccharum* | Sapindaceae | DTF | Deciduous |
| *Alnus acuminata* | Betulaceae | UMF | Evergreen |
| *Alnus rugosa* | Betulaceae | DTF | Deciduous |
| *Apeiba glabra* | Malvaceae | LTF | Deciduous |
| *Aspidosperma megalocarpon* | Apocynaceae | LTF | Evergreen |
| *Bellucia pentamera* | Melastomataceae | LTF | Evergreen |
| *Betula alleghaniensis* | Betulaceae | DTF | Deciduous |
| *Betula papyrifera* | Betulaceae | DTF | Deciduous |
| *Brosimum utile* | Moraceae | LTF | Evergreen |
| *Cariniana pyriformis* | Lecythidaceae | LTF | Deciduous |
| *Casearia arborea* | Salicaceae | LTF | Evergreen |
| *Cavendishia cordifolia* | Ericaceae | UMF | Evergreen |
| *Cecropia peltata* | Urticaceae | LTF | Evergreen |
| *Cedrela montana* | Meliaceae | UMF | Deciduous |
| *Cespedesia spathulata* | Ochnaceae | LTF | Deciduous |
| *Citharexylum dryanderae* | Verbenaceae | UMF | Evergreen |
| *Clathrotropis brachypetala* | Leguminasae | LTF | Evergreen |
| *Clusia multiflora* | Clusiaceae | UMF | Evergreen |
| *Cordia alliodora* | Boraginaceae | LTF | Deciduous |
| *Cordia cylindrostachya* | Boraginaceae | UMF | Deciduous |
| *Cordia spp.* | Boraginaceae | UMF | Deciduous |
| *Croton killipianus* | Euphorbiaceae | LTF | Deciduous |
| *Drimys granadensis* | Winteraceae | UMF | Evergreen |
| *Duguetia antioquensis* | Annonaceae | LTF | Evergreen |
| *Fagus grandifolia* | Fagaceae | DTF | Deciduous |
| *Fraxinus americana* | Oleaceae | DTF | Deciduous |
| *Goupia glabra* | Goupiaceae | LTF | Deciduous |
| *Hieronyma alchorneoides* | Phyllanthaceae | LTF | Evergreen |
| *Hymenaea courbaril* | Leguminosae | LTF | Evergreen |
| *Ilex nervosa* | Aquifoliaceae | UMF | Evergreen |
| *Iryanthera megistocarpa* | Myristicaceae | LTF | Evergreen |

| Species | Family | Biome | Leaf habit |
|---|---|---|---|
| *Juglans cinerea* | Juglandaceae | DTF | Deciduous |
| *Juglans neotropica* | Juglandaceae | UMF | Deciduous |
| *Lecythis ampla* | Lecythidaceae | LTF | Deciduous |
| *Miconia biappendiculata* | Melastomataceae | UMF | Evergreen |
| *Morella parvifolia* | Myricaceae | UMF | Deciduous |
| *Muntingia calabura* | Muntingiaceae | LTF | Evergreen |
| *Myrsine coriacea* | Primulaceae | UMF | Evergreen |
| *Ochoterenaea colombiana* | Anacardiaceae | LTF | Evergreen |
| *Ochroma pyramidale* | Malvaceae | LTF | Evergreen |
| *Oreopanax bogotensis* | Araliaceae | UMF | Deciduous |
| *Ostrya virginiana* | Betulaceae | DTF | Deciduous |
| *Piper bogotense* | Piperaceae | UMF | Evergreen |
| *Populus grandidentata* | Salicaceae | DTF | Deciduous |
| *Populus tremuloides* | Salicaceae | DTF | Deciduous |
| *Protium aracouchini* | Burseraceae | LTF | Evergreen |
| *Prunus buxifolia* | Rosaceae | UMF | Deciduous |
| *Prunus serotina* | Rosaceae | DTF | Deciduous |
| *Pseudoxandra sclerocarpa* | Annonaceae | LTF | Evergreen |
| *Quercus rubra* | Fagaceae | DTF | Deciduous |
| *Solanum humboldtianum* | Solanaceae | UMF | Evergreen |
| *Tabebuia guayacan* | Bignoniaceae | LTF | Deciduous |
| *Tapirira guianensis* | Anacardiaceae | LTF | Evergreen |
| *Tilia americana* | Malvaceae | DTF | Deciduous |
| *Trema micrantha* | Cannabaceae | LTF | Evergreen |
| *Ulmus americana* | Ulmaceae | DTF | Deciduous |
| *Verbesina crassiramea* | Compositae | UMF | Deciduous |
| *Viburnum lasiophyllum* | Adoxaceae | UMF | Evergreen |
| *Vismia macrophylla* | Hypericaceae | LTF | Evergreen |

**Table S3.** List of functional traits considered in this study with the abbreviations used in the text, the units of expression, and the hypothesized ecological function associated with the trait (Baraloto et al., 2010, Fortunel et al., 2012).

| Parameter | Abbreviation | Units | Ecological role |
|---|---|---|---|
| Leaf size | LS | $mm^2$ | Resource acquisition |
| Leaf thickness | LT | mm | Resource acquisition and defense |
| Leaf dry matter content | LDMC | $mg\ g^{-1}$ | Resource acquisition and defense |
| Specific leaf area | SLA | $mm^2\ mg^{-1}$ | Resource acquisition and defense |
| Photosynthetic capacity | Amass | $nmol\ CO_2\ g^{-1}\ s^{-1}$ | Resource acquisition |
| Foliar nitrogen | Leaf N | % | Resource acquisition and defense |
| Foliar phosphorus | Leaf P | $mg\ kg^{-1}$ | Resource acquisition |
| Foliar calcium | Leaf Ca | $mg\ kg^{-1}$ | Resource defense |
| Foliar magnesium | Leaf Mg | $mg\ kg^{-1}$ | Resource acquisition and defense |
| Stem density | SD | $mg\ mm^{-3}$ | Hydraulic transport, mechanical strength and defence |
| Branch density | BD | $mg\ mm^{-3}$ | Hydraulic transport, mechanical strength and defence |
| Tree height | H | m | Resource capture and reproduction |

**Table S4.** Summary of phylogenetic hierarchical Bayesian models that examine the variation in NSC concentrations of roots, stems, branches and leaves.

| | **Roots** | | **Stems** | | **Branches** | | **Leaves** | |
|---|---|---|---|---|---|---|---|---|
| Fixed effect | Estimate | Effective sample size | Estimate | Effective sample size | Estimate | Effective sample size | Estimate | Effective sample size |
| Intercept | 10.67 (7.11 - 14.62) | 7098 | 0.32 (-0.19 - 0.9) | 9442 | 2.05 (1.56 - 2.55) | 9379 | 8.35 (7.05 - 9.71) | 7268 |
| Biome LTF | 0.49 (-3.92 - 4.93) | 5307 | 0.84 (0.19 - 1.48) | 10439 | -0.41 (-0.78 - -0.04) | 9967 | -4.31 (-5.88 - -2.73) | 6724 |
| Biome UMF | 0.55 (-4.2 - 5.19) | 5192 | 1.12 (0.42 - 1.8) | 10001 | 0.01 (-0.37 - 0.39) | 9740 | -3.02 (-4.71 - -1.3) | 6776 |
| Leaf habit (Evergreen) | -2.63 (-6.05 - 0.82) | 8131 | -0.03 (-0.52 - 0.46) | 11222 | -0.22 (-0.52 - 0.07) | 11598 | 0.13 (-1.04 - 1.28) | 9677 |
| Fast-slow PC1 | -0.3 (-1.7 - 1.01) | 5986 | 0.1 (-0.08 - 0.28) | 16680 | -0.01 (-0.12 - 0.1) | 13431 | -1.11 (-2.24 - 0.04) | 7861 |
| Fast-slow PC2 | 1.4 (-0.21 - 2.97) | 9429 | 0.08 (-0.18 - 0.34) | 10869 | -0.03 (-0.17 - 0.12) | 12712 | 0.52 (-0.12 - 1.15) | 10094 |
| LTF x Fast-slow PC1 | | | | | | | 1.71 (0.35 - 3.05) | 9420 |
| UMF x Fast-slow PC1 | | | | | | | 1.43 (0.12 - 2.73) | 8172 |

The effective sample size reports the number of independent samples with the same estimation power as the N autocorrelated samples (Kruschke, 2015).

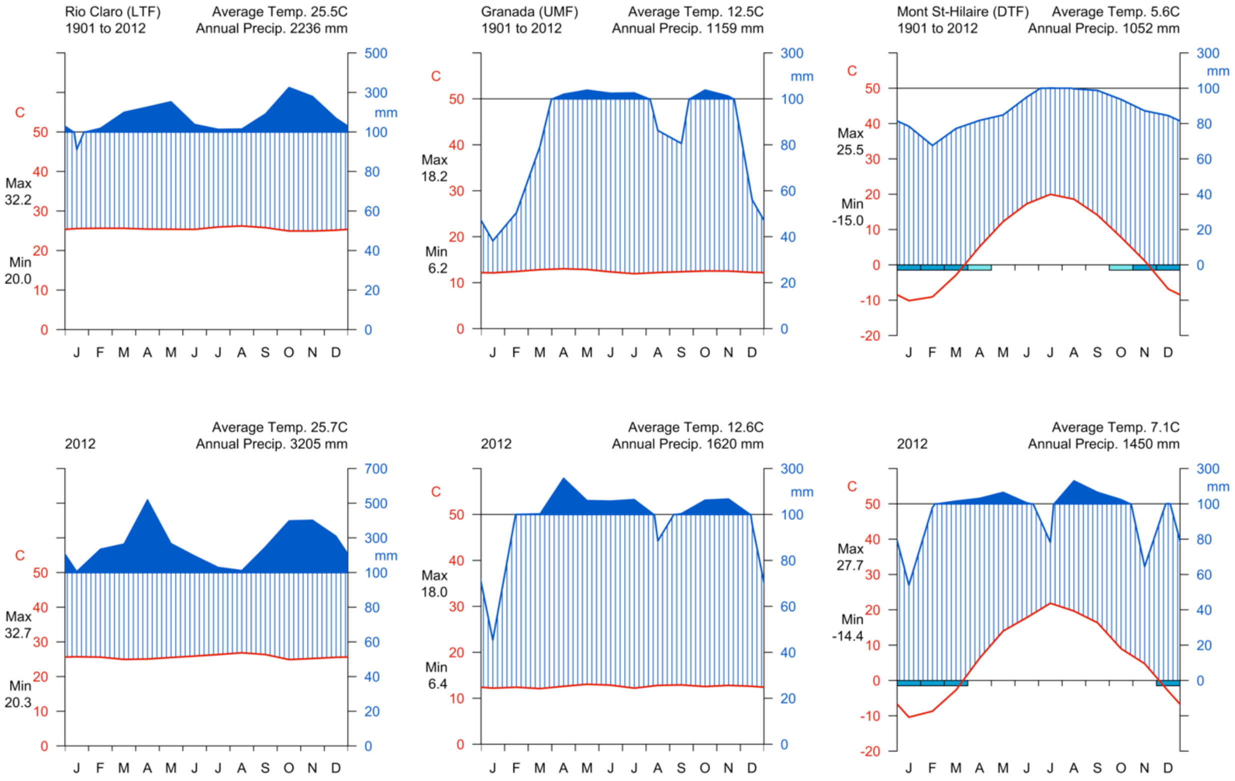


**Fig. S1.** Walter-Lieth climatic diagram of the three study sites. Data from high-resolution gridded dataset of the Climatic Research Unit (CRU) at the University of East Anglia (Harris et al., 2014). Above, average climate data from 1901 to 2012 and below climate from 2012. UMF: upper montane tropical forest, LTF: lowland tropical rainforest, and DTF: deciduous temperate forest.

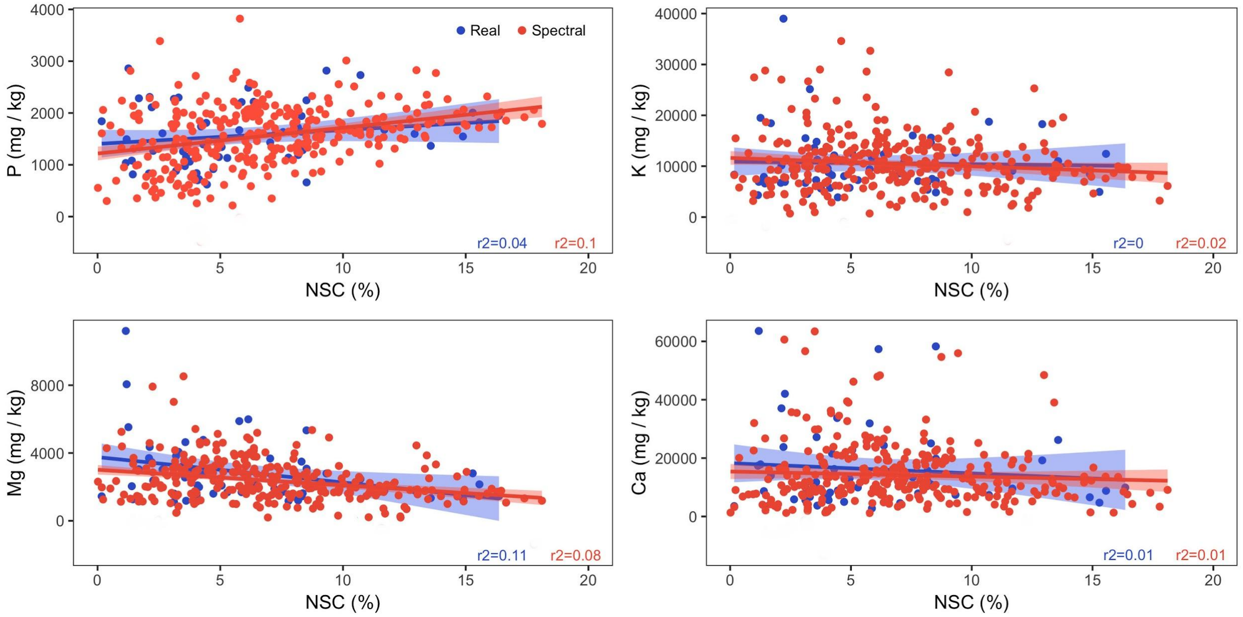


**Fig. S2.** Relationships between direct measures and imputed spectral values of leaf NSC and functional traits. Refer to Table S3 for trait abbreviations.

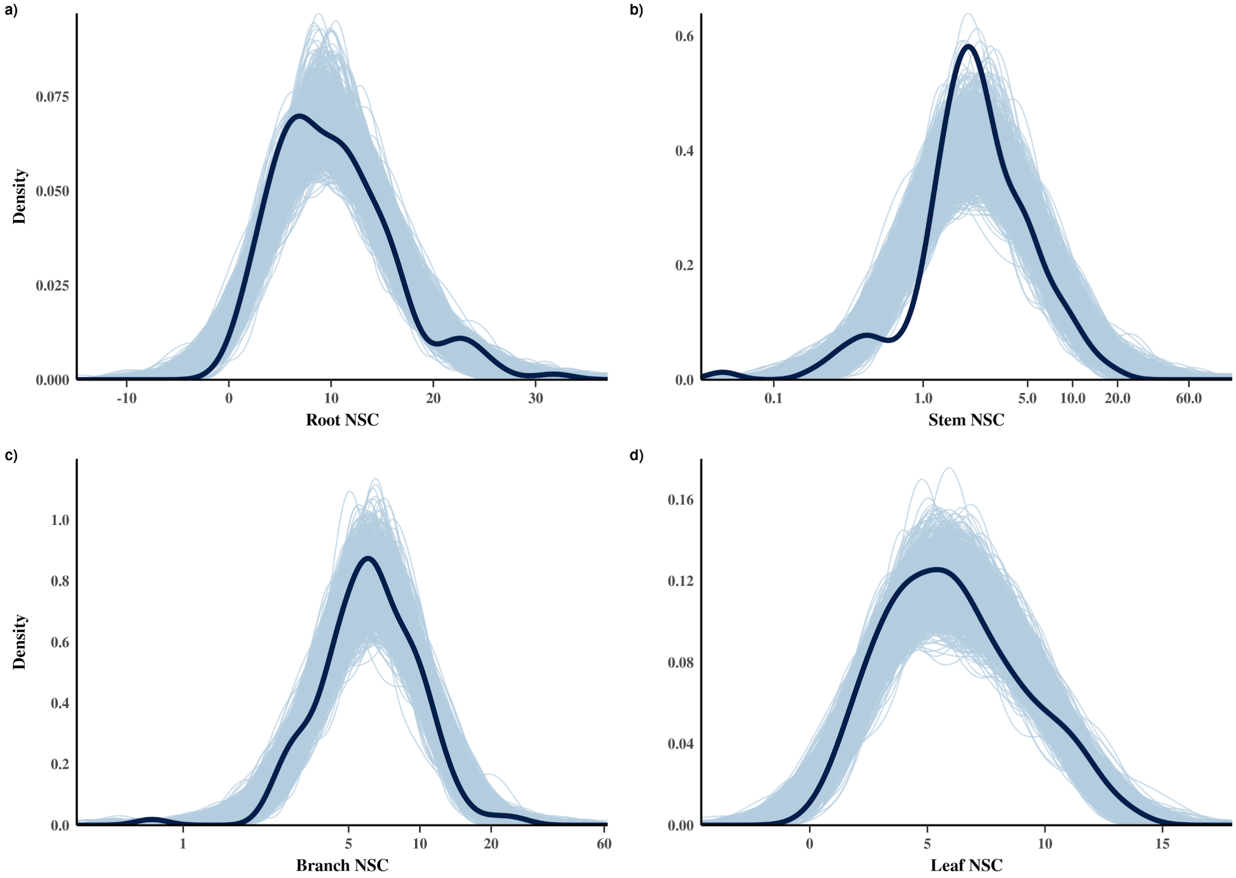


**Fig. S3.** Posterior predictive distributions for phylogenetic hierarchical Bayesian models that estimate variation in NSC concentrations of **a**) roots, **b**) stems, **c**) branches, and **d**) leaves. Dark blue lines represent the observed data, while the light blue lines represent simulations of the response variable (*n* = 1,000 posterior samples).

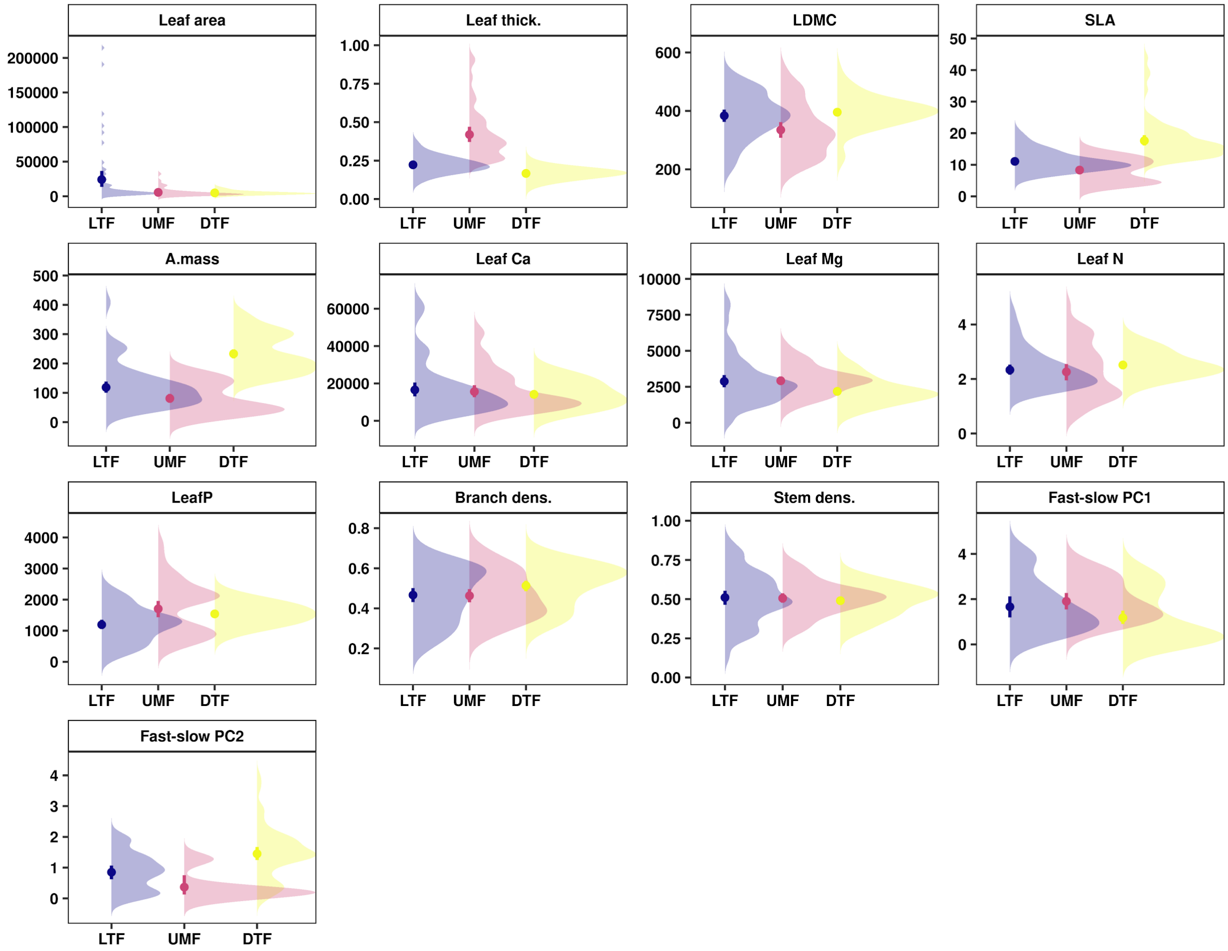


**Fig. S4.** Trait means and distributions of plant functional traits and fast-slow PCs across biomes. Whisker bars show means and 95% confidence intervals. LTF: lowland tropical rainforest, UMF: upper montane forest, and DTF: deciduous temperate forest. Table S3 shows trait abbreviations.

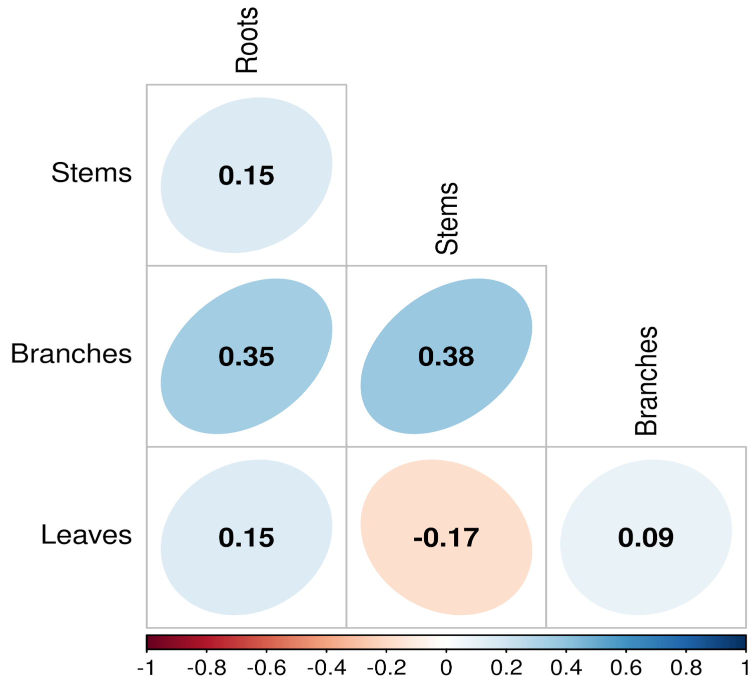


**Fig. S5.** Pairwise correlations of NSC carbohydrates among tree organs across three biomes.

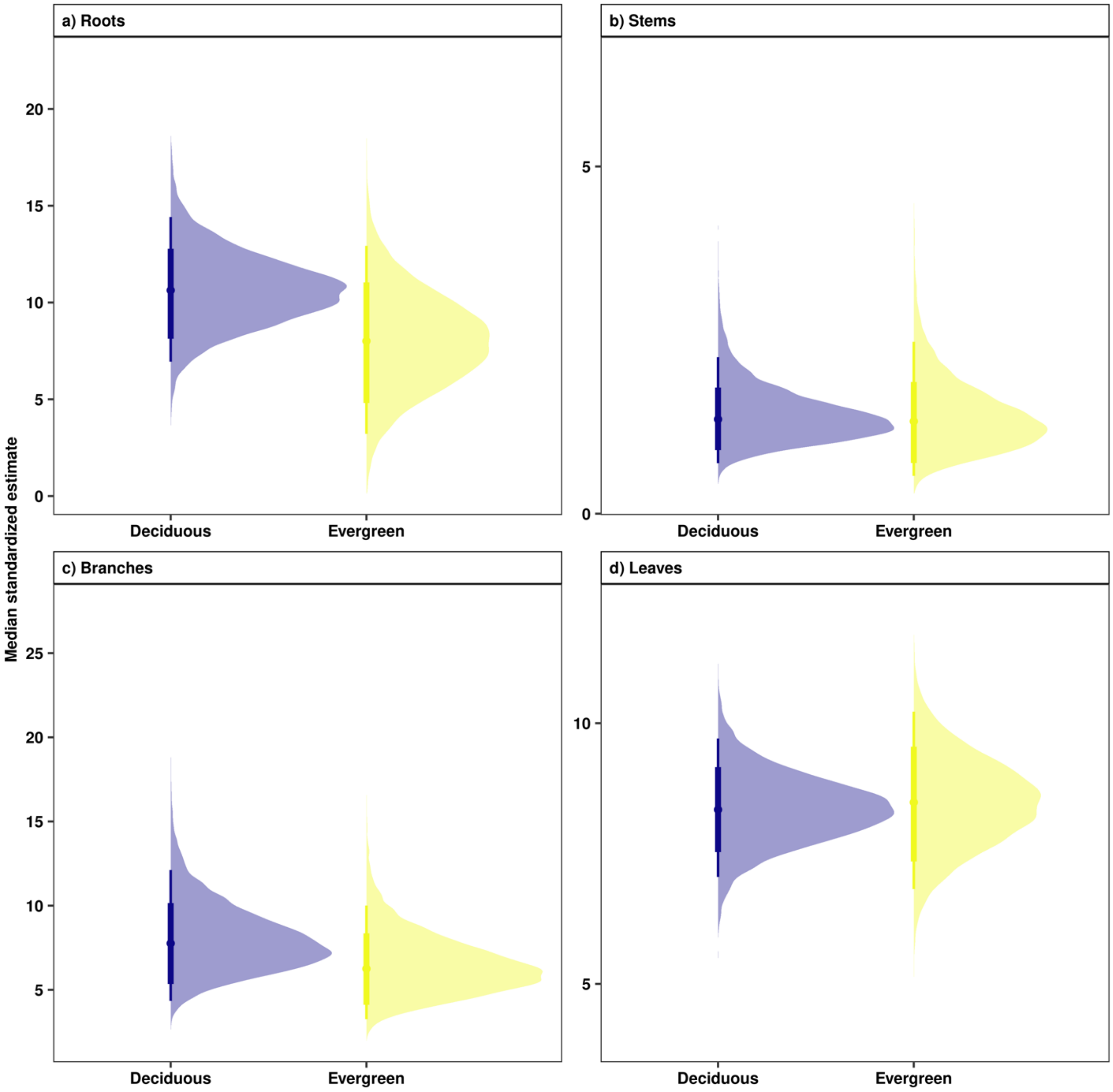


**Fig. S6.** Model estimates of NSC concentrations for deciduous and evergreen species for **a**) roots, **b**) stems, **c**) branches, and **d**) leaves. Points are medians and whisker bars are 80% and 95% credible intervals, which were estimated using phylogenetic hierarchical Bayesian models.

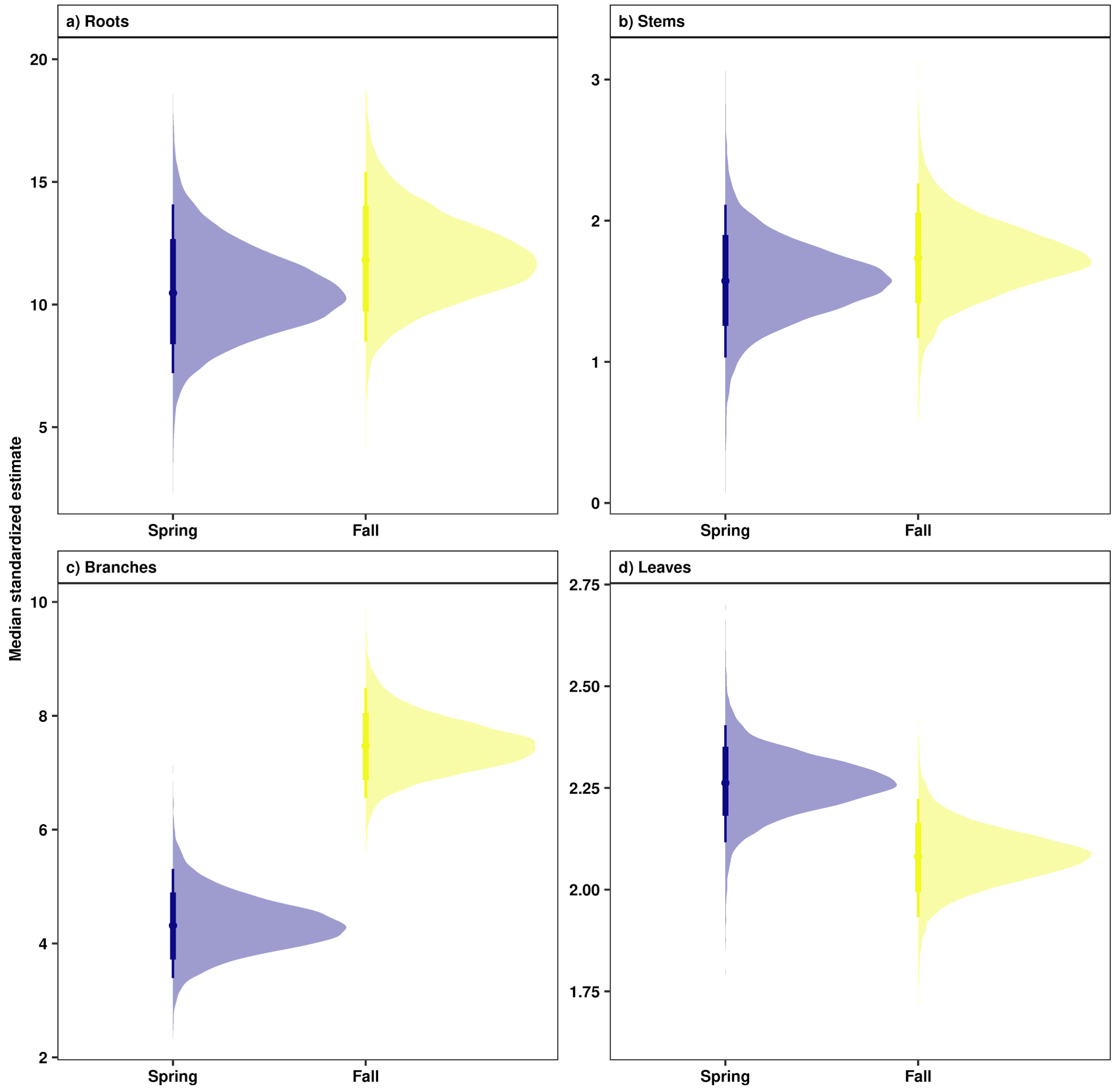


**Fig. S7**. Model estimates of NSC concentrations for **a**) roots, **b**) stems, **c**) branches, and **d**) leaves across seasons in a deciduous temperate forest. Points are medians and whisker bars are 80% and 95% credible intervals, which were estimated using phylogenetic hierarchical Bayesian models.

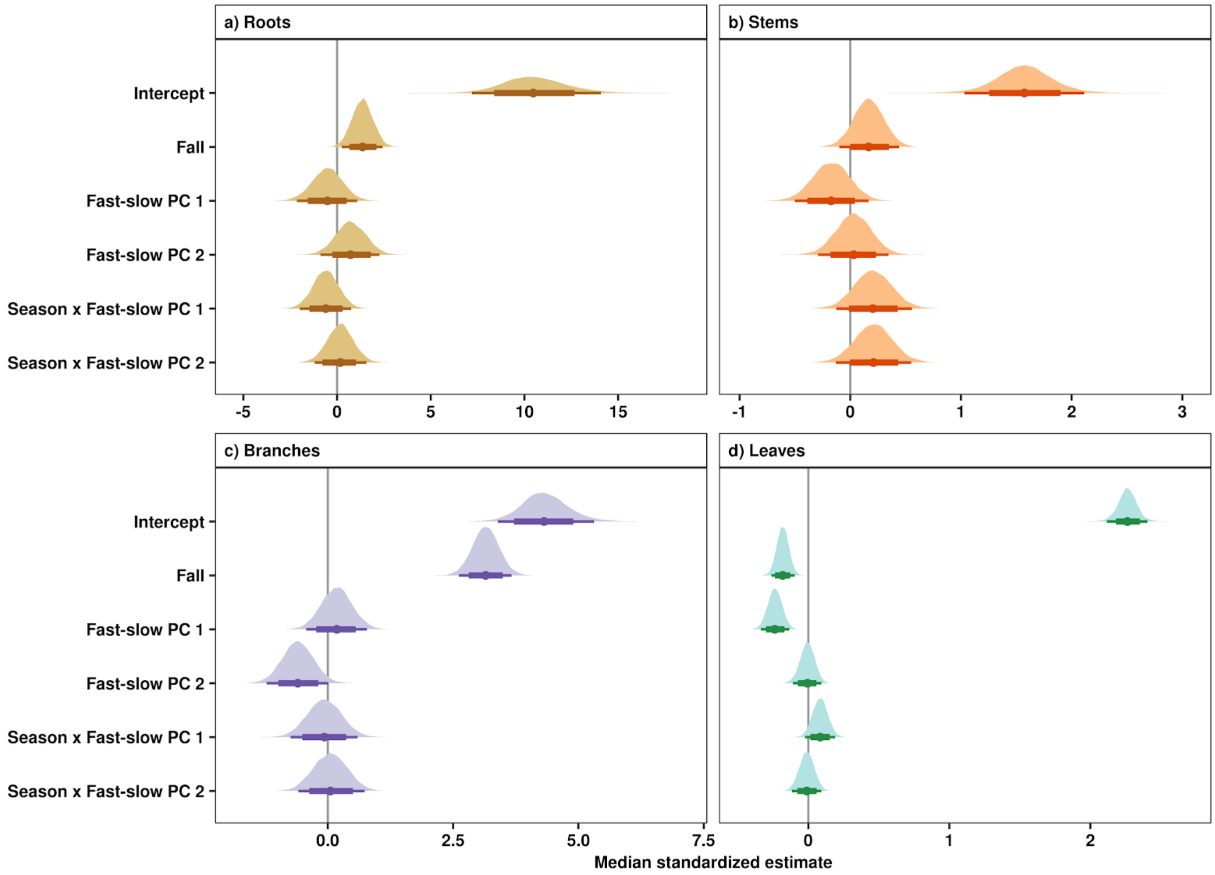


**Fig. S8**. Influence of the fast-slow continuum and seasons on NSC concentrations of deciduous temperate forest (DTF) in **a**) roots, **b**) stems, **c**) branches, and **d**) leaves. Points are medians and whisker bars are 80% and 95% credible intervals and were estimated using phylogenetic hierarchical Bayesian models. Continuous variables were z-transformed prior to analysis to facilitate comparisons (within and across tree organs). Fast-slow PC1 and fast-slow PC2 are the first two axes of a principal component analysis of fast-slow plant functional traits.
